# A straightforward approach for bioorthogonal labeling of proteins and organelles in live mammalian cells, using a short peptide tag

**DOI:** 10.1101/708545

**Authors:** Inbar Segal, Dikla Nachmias, Eyal Arbely, Natalie Elia

## Abstract

In the high-resolution microscopy era, genetic code expansion (GCE)-based bioorthogonal labeling offers an elegant way for direct labeling of proteins in live cells with fluorescent dyes. This labeling approach is currently not broadly used live cell applications, partly because it needs to be adjusted to the specific protein under study. Here, we present a generic, 14-residues long, N-terminal tag for GCE-based labeling of proteins in live mammalian cells. Using this tag, we generated a library of GCE-based organelle markers, demonstrating the applicability of the tag for labeling a plethora of proteins and organelles. Finally, we show that the HA epitope, used as a backbone in our tag, can be substituted with other epitopes and, in some cases, can be completely removed, reducing the tag length to 5 residues. The GCE-tag presented here offers a powerful, easy-to-implement tool for live cell labeling of cellular proteins with small and bright probes.

## BACKGROUND

Tracking the dynamics of proteins and organelles in live cells is key to understanding their functions. For this, fluorescent protein (e.g. GFP) or self-labeling protein (e.g. Halo-Tag) tags are routinely attached to proteins in cells (1). While these tags are vigorous and easy-to-implement, they are large and bulky (e.g. GFP, ~27 kDa; Halo-tag, 33 kDa), such that their attachment could affect the dynamics and function of the protein under study. Using genetic code expansion (GCE) and bioorthogonal chemistry, it is now possible to non-invasively attach fluorescent dyes (Fl-dyes) to specific protein residues, thereby allowing essentially “tag-free” labeling of proteins in live cells (1–3). Indeed, this approach has been applied, in recent years, for fluorescent labeling of extra and intra cellular proteins (4–10).

In GCE-based labeling, a non-canonical amino acid (ncAA) carrying a functional group is incorporated into the sequence of a protein in response to an in-frame amber stop codon (TAG), via an orthogonal tRNA/tRNA-synthetase pair (reviewed in (11, 12)). Labeling is then carried out by a rapid and specific bioorthogonal reaction between the functional group and the Fl-dye (2, 4, 8, 9, 13, 14). Successful GCE-based labeling hence relies on exogenous expression of an orthogonal tRNA/tRNA-synthetase pair and a protein of interest (bearing a ncAA) at sufficient levels to allow effective labeling.

The ncAA (and consequently the Fl-dye) can, in theory, be incorporated anywhere in the protein sequence. In practice, however, finding a suitable labeling site can be laborious and time-consuming for several reasons. First, prior knowledge or functional assays are necessary to ensure that insertion of the ncAA at a specific position does not affect protein structure and function (4–7, 10). Second, the efficiency of ncAA incorporation varies at different locations in the protein with no guidelines for the preferred sequence context having been reported (3–7, 15). Notably, low efficiency of ncAA incorporation does not only lead to ineffective labeling but also to translation of a truncated version of the protein (resulting from the insertion of a premature stop codon), which can be toxic to cells (5, 6, 16, 17). Third, the ncAA should be incorporated in a position that will allow the functional group to be accessible to the solvent in order to enable efficient bioorthogonal conjugation with the Fl-dye. All these requirements are protein-specific, such that any attempt at labeling via this approach begins with a screen for suitable incorporation sites (2-4, 6, 7). Consequently, despite its great potential, GCE-based labeling is presently not widely used in mammalian live cell imaging studies (5).

To bypass the screening steps currently associated with GCE-based labeling and expand the use of the technique, we set out to design a minimal peptide tag that encodes the incorporation a ncAA through GCE for bioorthogonal labeling of proteins in live cells; we term this tag a GCE-tag (Fig. 1a). Using the HA epitope as a backbone we optimized a 14-residue long N terminal tag and demonstrated its applicability for live cell labeling of a variety of proteins and organelle tags in mammalian cells. With this tag in hands, essentially any protein in cell can now be labeled via GCE and bioorthogonal chemistry for live cell imaging applications with no prior knowledge or screening steps required.

**Fig 1.**
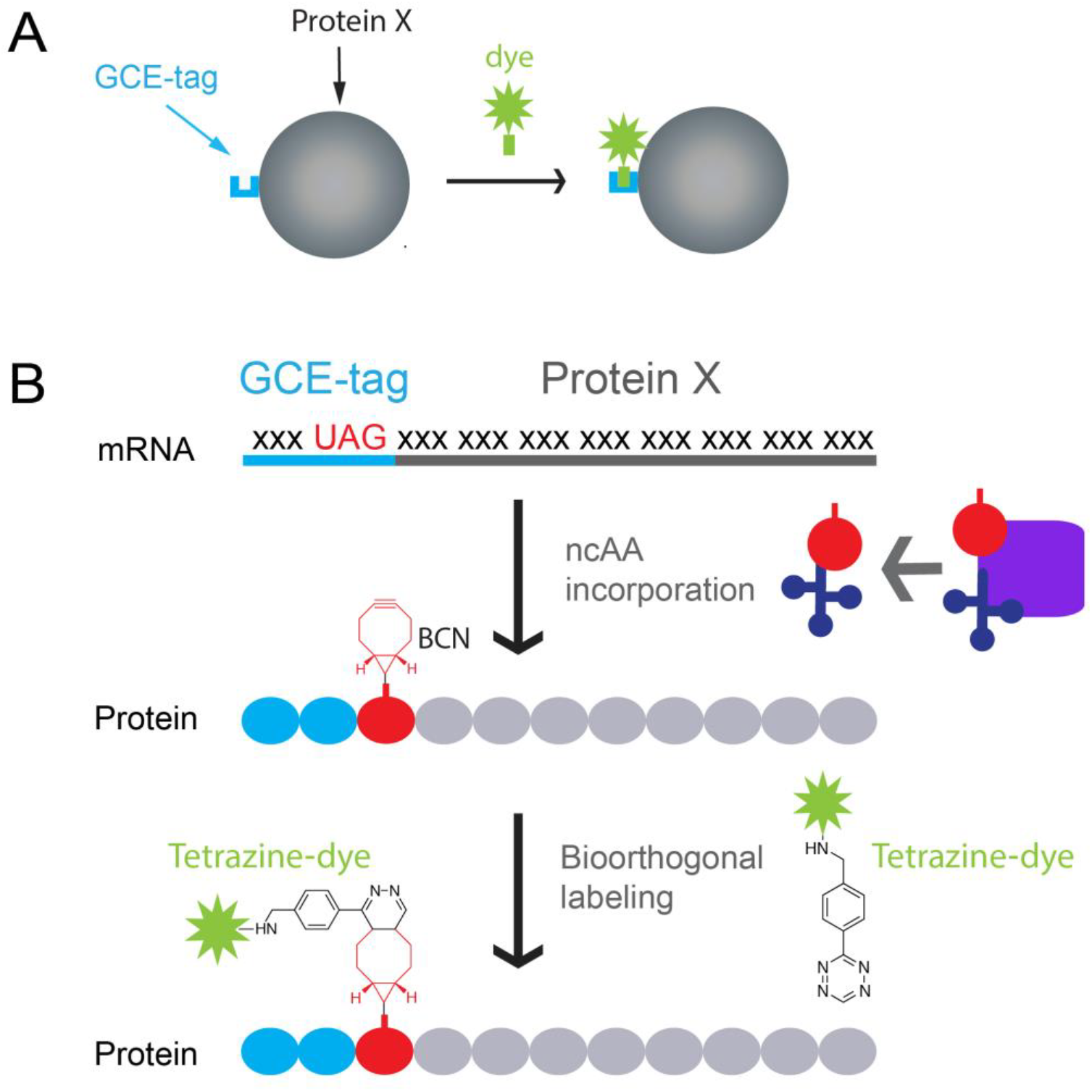
Using a GCE-tag for fluorescence labeling of proteins via genetic code expansion (GCE) and bioorthogonal chemistry. (A) Schematic representation of the labeling approach. A GCE-tag (blue) is attached to the N terminal of the protein (grey). Binding of the Fl-dye (green) to the GCE-tag results in labeling of the protein. (B) Graphical illustration of the experimental design. The stop codon, UAG, and nucleotides encoding a short polypeptide tag are added upstream to the coding sequence of a protein. During ribosomal translation the ncAA BCN-Lysine (red) is incorporated into the protein in response to the in-frame UAG codon using a specific, orthogonal pair of tRNA (dark blue) / tRNA-Synth (purple). Finally, a tetrazine–conjugated Fl-dye (green) is covalently attached to BCN-Lysine via a bioorthogonal reaction. As a result, the protein is directly labeled with Fl-dye via a small polypeptide tag.

## RESULTS

Proteins can potentially be tagged at their N or C terminal. For GCE-based labeling, we chose to design an N terminal tag in order to avoid protein truncations resulting from inefficient incorporation of the ncAA (5, 6, 16, 17). On the basis of our previous work, potential tags were cloned into a single expression vector, designed to encode the incorporation of the ncAA bicyclo-nonyne Lysine (BCN-Lysine) that bioorthogonally reacts with Tetrazine conjugated Fl-dyes (Fig. 1b and Figure S1A) (18). Labeling potency was initially assessed using α-tubulin as a benchmark and evaluating microtubule (MT) labeling in live mammalian cells in the presence of tetrazine-silicon-rhodamine (SiR-Tet) (6, 19). MT labeling obtained upon site-specific incorporation of BCN-Lysine at α-tubulin position 45 (α-tubulin^45TAG^) was used as a reference, given our earlier demonstration of the efficacy of MT labeling at this site (6).

The most minimal N-terminal GCE-tag should include a TAG codon that will encode the incorporation of BCN-Lys. Unfortunately, MT labeling was not obtained upon expressing α-tubulin that carries a TAG codon at its N terminal (directly before the first residue) in the presence of SiR-Tet (Figure S1B and S2). This indicated that the sole incorporation of BCN-Lys at the N terminal of α-tubulin is not sufficient for labeling. Potential N-terminal tags, bearing different length, were therefore designed based on the commonly used 9 amino-acid (AA)-long hemagglutinin (HA) epitope as backbone (20) (Fig. 2a and Figure S1C-F). The TAG codon was either introduced within the HA sequence (replacing the last codon in the epitope) (1), immediately after the HA sequence (2) or after a short commonly used flexible linker comprising either glycine-serine (GS; 3) or glycine-glycine-serine-glycine (GGSG; 4). Low but noticeable levels of α-tubulin were observed using any of these tags, indicating that N-terminally tagged α-tubulin was expressed in cells (Fig. 2b). In live cell labeling experiments with SiR-Tet, little to no MT decoration was obtained using α-tubulin tagged with probes 1 or 2 (Fig. 2c, 1-2), while clear and specific labeling was observed using tags 3 and 4 (Fig. 2c, 3-4). Signal-to-noise ratios (SNRs) were significantly higher in cells expressing α-tubulin tagged with probe 4 (as compared to probe 3) and even higher than those obtained in cells expressing α-tubulin^45TAG^ (Fig. 2d, e). We, therefore, concluded that N’ HA-GGSG is the minimal GCE-tag suitable for α-tubulin labeling in live mammalian cells.

**Fig 2.**
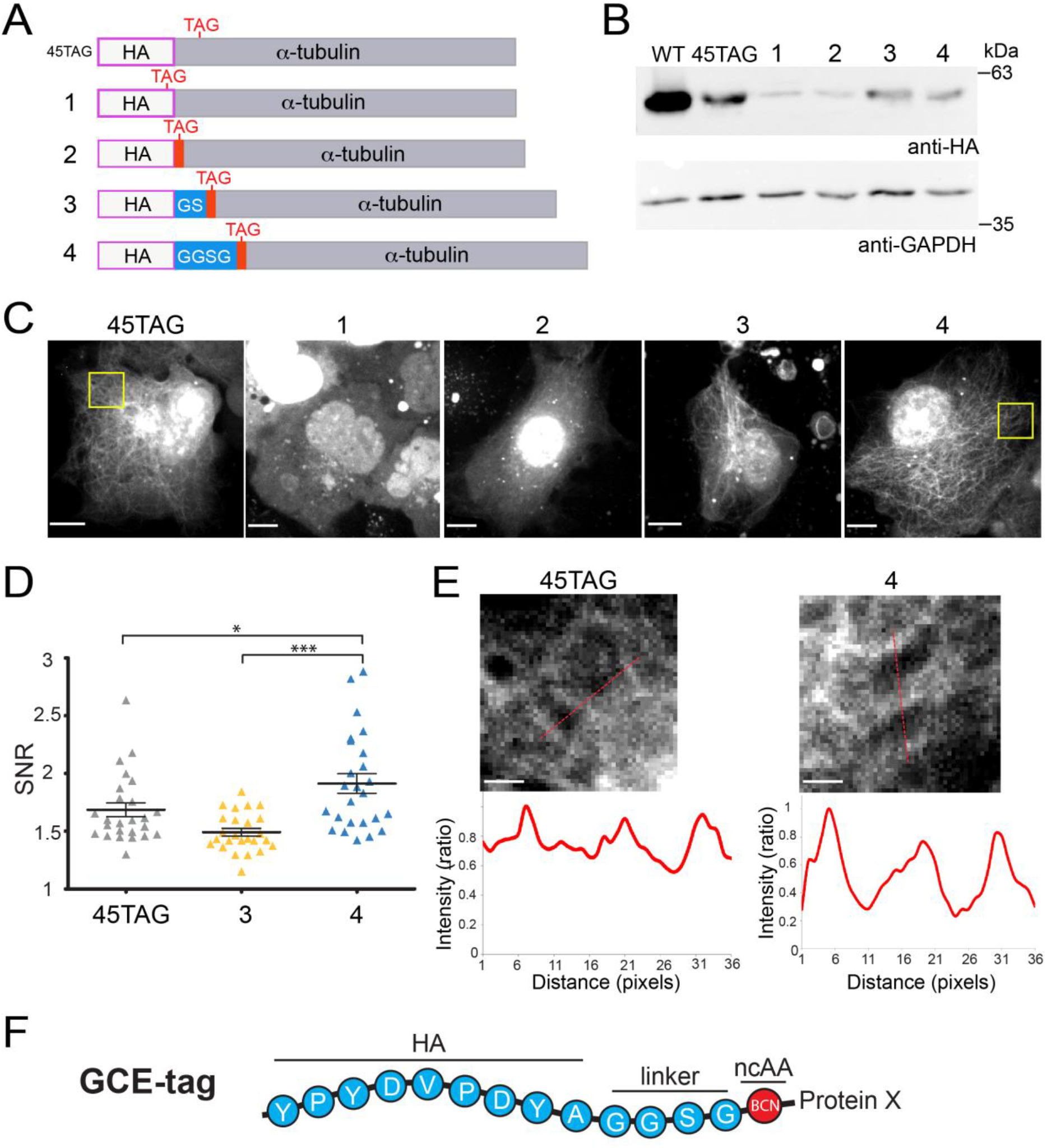
Optimizing a minimal tag for genetic code expansion (GCE)-based fluorescence labeling. (A) Schematic view of the tags used and tested in (B-D). Complete sequences of the tags are shown in Figure S1C-F. (B-E). GCE-tags (specified in A) were sub-cloned to the N terminal of α-tubulin using the pBUD-Pyl-RS-tub plasmid (Figure S1A). HEK293T (B) or COS7 (C-E) cells were transfected with pBUD-Pyl-RS-tub plasmids carrying different tags and incubated for 48h in the presence of the ncAA BCN-Lysine. Cells were then subjected to western blot analysis (B) or labeled with SiR-tet for 1h and imaged live (C-E). Images in C represent maximum intensities of 3D z-stacks of representative cells. (D) SNR values measured in cells expressing α-tubulin with BCN-Lysine incorporated either in position G45 or in the N-terminal via tags 3 or 4. Statistical significance was determined by ANOVA ***p<0.0001, * p<0.05 (n=25). (E) Zoomed-in images of the regions marked in yellow squares in e. Intensity values along the red line drawn in the images are plotted below, demonstrating the improvement in SNR. Scale-bar (C): 10 μm, (E) 2 μm. Results presented (B-E) were obtained in at least 3 independent experiments. (F) Schematic view of the optimized GCE-tag used for further labeling.

To test the applicability of the GCE-tag (HA-GGSG, Fig. 2f and Figure S1F) for labeling diverse cellular compartments, we cloned the sequences of known organelle markers into our GCE single expression vector system, in-frame with the GCE-tag. Specifically, we used GFP-CAAX as a plasma membrane (PM) marker, GFP-SKL as a peroxisomal marker, Lamp1 as a lysosomal marker, CD63 as a marker of multivesicular bodies (MVBs), ER^cb5^TM as a endoplasmic reticulum (ER) marker, Exo70 as an exosomal marker and Mito^cb5^TM(mito) or mito-DsRed as mitochondrial markers (see Table S1 for sequences). Nuclear labeling was avoided due to the known non-specific labeling of the nucleus observed with GCE-based labeling (Fig. 2c) (4, 6, 16).

Clear PM labeling was obtained in both the GFP and SiR channels upon expressing GCE-tagged GFP-CAAX. Moreover, SNRs measured for PM labeling were similar using either GCE-tagged GFP-CAAX or the optimized ncAA incorporation site in GFP (21) (compare upper and lower panels in Fig. 3a). In COS7 cells expressing the peroxisome marker GFP-SKL fused to the GCE-tag and labeled with SiR-Tet, remarkable co-localization of GFP and SiR in small puncta was observed throughout the cell (Fig. 3b). Apart from the non-specific nuclear labeling obtained in the SiR channel, essentially all SiR-Tet-labeled puncta were positive for GFP (Pearson correlation = 0.814). Together, these data indicate that both the GCE-tagged PM and peroxisome markers are expressed and properly targeted in cells.

**Fig 3.**
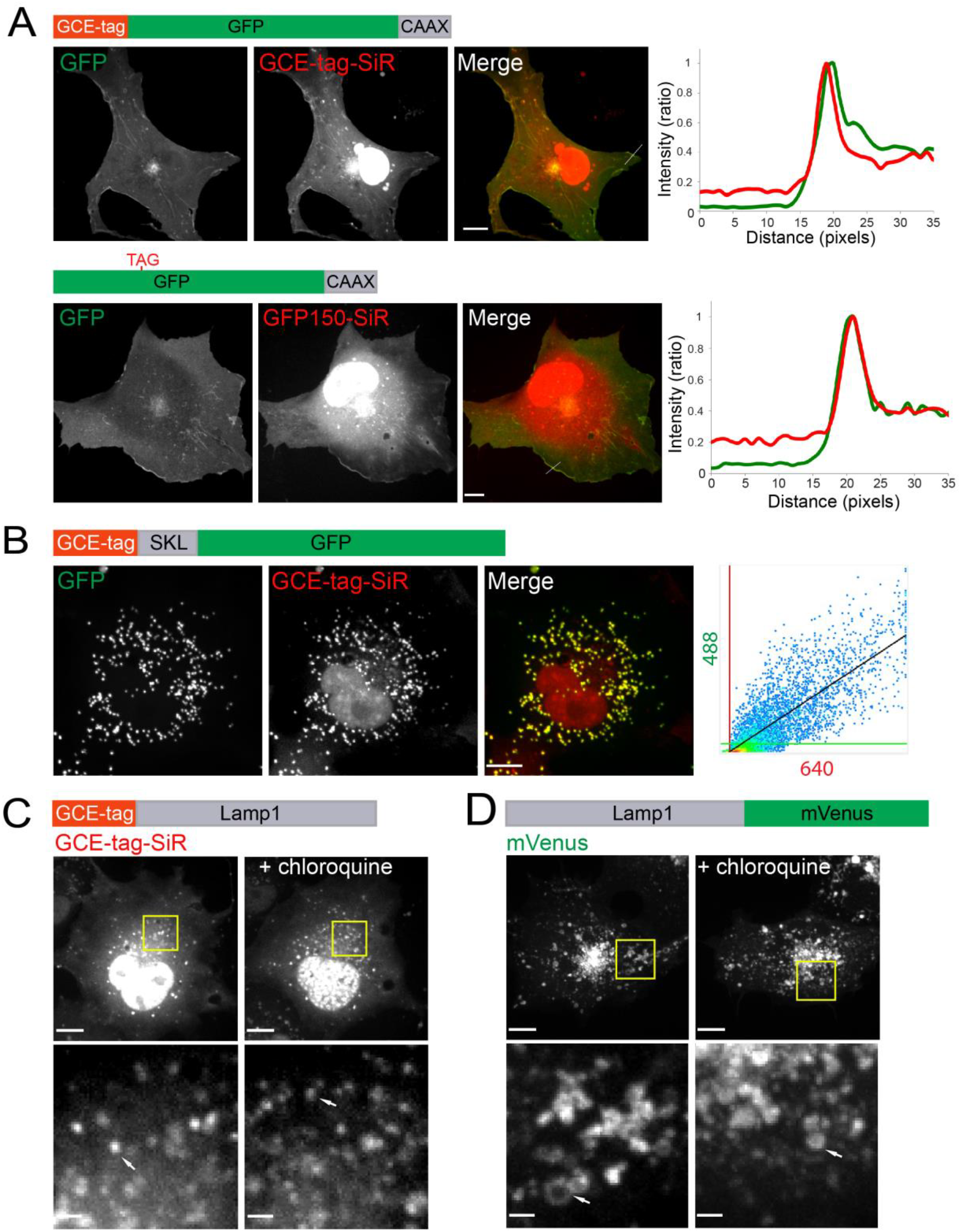
Plasma membrane (PM), peroxisomes and lysosomes are efficiently labeled in live cells via the minimal GCE-tag and Tetrazine-SiR. Live cell images of COS7 cells expressing different organelle markers tagged with the GCE-tag (C), a fluorescent protein (D) or both (A and B). Complete sequences are provided in Table S1. Cells labeled bioorthogonally were transfected with pBUD-Pyl-RS plasmids containing different organelle protein markers (indicated below) with GCE-tag fused to their N terminal. Cells were then incubated for 48 h in the presence of BCN-Lysine, labeled with SiR-Tet and imaged. (A) Cells expressing GCE-tag-GFP-CAAX (upper panel) or GFP-CAAX mutated at position 150 to incorporate the ncAA BCN-Lysine (bottom panel) and labeled with SiR-Tet showing plasma membrane labeling. Normalized intensity values of GFP and SiR-Tet along the red lines drawn in merged images are shown in the plots to the right. Intensity values were normalized to background levels. (B) Peroxisome labeling in cells expressing GCE-tag-GFP-SKL. Plot on the right represents colocalization analysis between intensity values of the 488 (GFP) and 640 (SiR) performed on a large subset of the cell that excludes the nucleus, Pearson correlation value = 0.814. (C-D) Cells expressing GCE-tag-Lamp1 labeled with SiR-Tet (C) or Lamp1-mVenus (D). Left panels: no treatment. Right panels: treatment with the lysosome inhibitor chloroquine (3 h, 120 μM). Zoomed in images of a subset of the cells (correspond to yellow squares in upper panels) are shown in bottom panels. Results presented in each panel were obtained in at least 3 independent experiments. Scale-bars: 10 μm, Zoomed-in images, 2 μm.

Effective lysosome labeling was obtained using GCE-tagged Lamp1 in the presence of SiR-Tet (Fig. 3c, d). Overall, labeling was comparable to that obtained with Lamp1-mVenus, with the expected increase in lysosome numbers upon chloroquine treatment being observed using either tag (i.e., GCE-tag or mVenus) (22). We noticed that while lysosome labeling using Lamp1-mVenus highlighted both filled and hollow structures, only filled structures were observed in cells labeled with GCE-tag-SiR-Lamp1. This difference may result from the position of the tag; the mVenus tag faces the cytosol, while the GCE-tag faces the lysosomal lumen. Labeling lysosomal lumen proteins using conventional fluorescent proteins is challenging due to the pH sensitivity of the latter (23). SiR florescence, on the other hand, is much less sensitive to pH (Figure S3) and is, therefore, suitable for intra-lysosome labeling.

Initial attempts at labeling MVBs, ER, mitochondria and exosomes using the GCE-tag and SiR-Tet resulted in no specific staining (Figure S4). Yet, in immunostaining experiments using anti-CD63 antibodies, GCE-tagged CD63 co-localized with endogenous CD63 (Fig. 4a). We thus reasoned that GCE-tagged CD63 was properly expressed and targeted in cells, yet failed to bind the Fl-dye via the bioorthogal reaction. Consistent with this notion, substituting TAMRA-Tet for SiR-Tet resulted in specific labeling of MVBs, ER and exosomes using GCE-tagged CD63, ER^cb5^TM and Exo70, respectively (Fig 4b, c and e). Mitochondrial labeling was not obtained using either SiR-or TAMRA-Tet together with the GCE-tag-Mito^cb5^TM or mito-DsRed probes (Figure S4B, C). The expression patterns observed with the GCE-tagged organelle markers that were successfully labeled (i.e., for MVBs, ER and exosomes) were similar to those obtained using conventional Fl-protein markers, validating labeling specificity (Fig. 4d, f).

**Fig 4.**
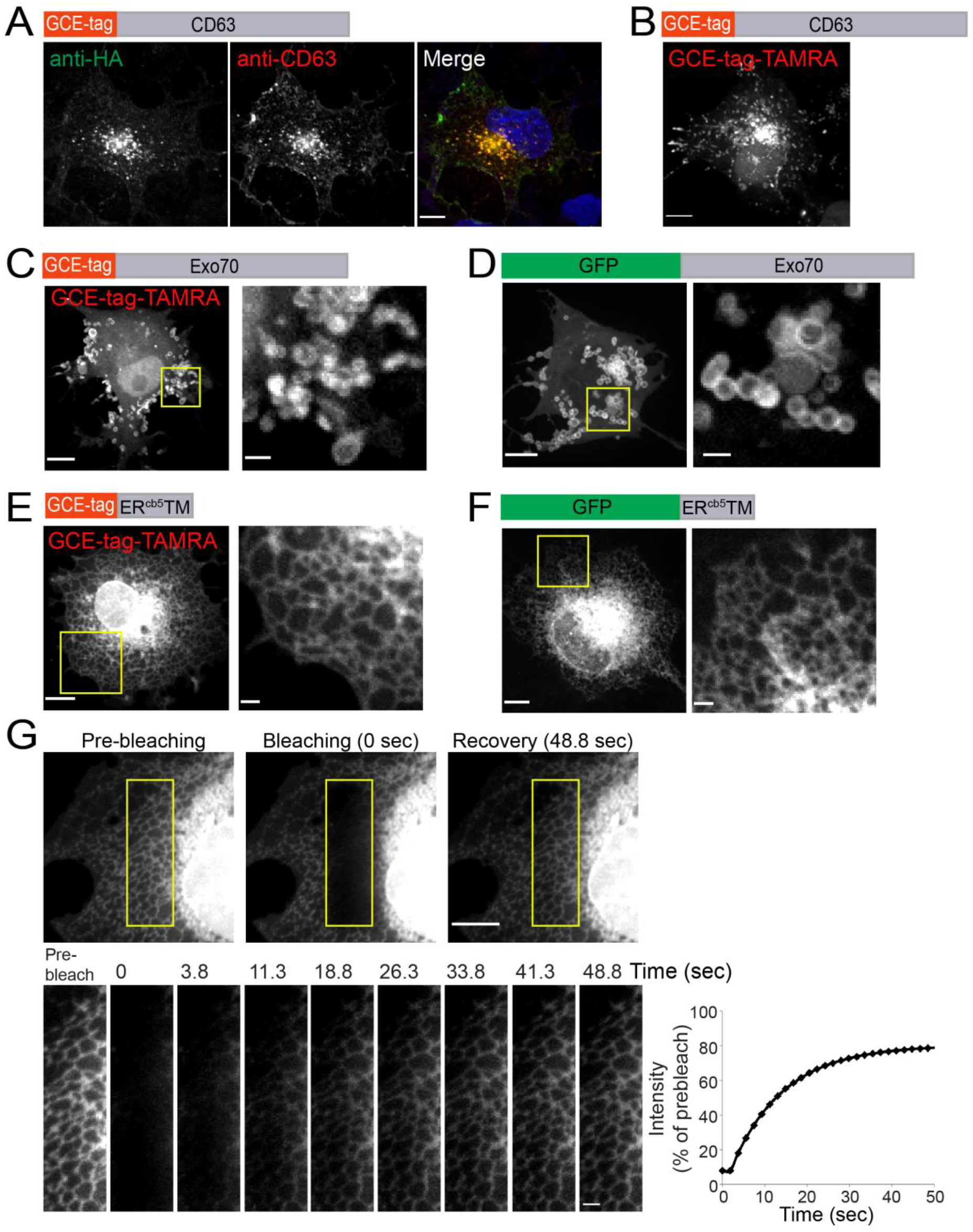
MVBs, exosomes and ER exhibit specific cellular labeling via the minimal GCE-tag in the presence of Tetrazine-TAMRA. (A) COS7 cells expressing the MVB marker CD63 conjugated to the GCE-tag were fixed and stained with anti-HA and anti-CD63 antibodies. (B) MVB labeling in cells expressing GCE-tag-CD63 labeled with TAMRA-Tet. (C and D) Exosome labeling in cells expressing GCE-tag-Exo70 labeled with TAMRA-Tet (C) or Exo70-GFP (D). (E-G) ER labeling in cells expressing GCE-tag-ER^cb5^TM labeled with TAMRA-Tet (E) or ER^cb5^TM-GFP (F). Right panels in (C-F) are zoomed in images of a subset of the cells (correspond to yellow squares in left panels). (G) FRAP analysis of cells expressing GCE-tag-ER^cb5^TM labeled with TAMTA-Tet. Snapshots of a subset of a representative cell taken from the movie sequence (Video S7) are shown in the upper panel. Sequential images of the bleached ROI are shown in bottom panel (Video S8). Plot on the right represents the exponential fit of fluorescence intensity recovery versus time after photobleaching in the ROI. Intensity values were corrected for unintentional bleaching and normalized to intensity levels measured pre-bleaching. Results presented in each panel were obtained in at least 3 independent experiments. Scale-bars: 10 μm, Zoomed-in images, 2 μm.

Fl-dye labeling of the GCE-tagged constructs allowed live cell recordings and tracking of any of the labeled cellular structures (i.e., MT, PM, peroxisomes, ER, MVB and exosomes; Videos S1-S8). Fluorescence recovery after photobleaching (FRAP) experiments were successfully performed using the GCE-ER^cb5^TM ER marker, with recovery times being comparable to those measured in the ER using VSVG-GFP (24) (Fig. 4g and Videos S7-S8). It is worth mentioning that the overall size of the GCE-tag-ER^cb5^TM ER marker is considerably smaller than VSVG-GFP (GCE-tag-ER^cb5^TM, ~8.5 kDa; VSVG-GFP, ~84.5 kDa). Taken together, these results indicate that the newly developed GCE-based organelle markers reported here can be employed to study organelle dynamics in live cells with much smaller probes.

The 14 residue-long GCE-tag described here comprises the complete sequence of the HA tag. As such, the HA epitope can still be exploited for other applications, such as immunoblot, immunofluorescence and immunoprecipitation (Fig. 2b, 4a and 5a). To expand the versatility of the tag, we tested whether the HA epitope can be replaced by the Myc (10 residues) or FLAG (8 residues) epitopes (Fig. 5a-d and Figure S5). GFP-SKL carrying HA/FLAG/Myc GCE-tags were successfully expressed in cells, with SiR-Tet fluorescence co-localizing with GFP (Pearson correlation values: FLAG, 0.858, Myc, 0.73) (Fig. 5a, b and Figure S5A). Efficient labeling with the FLAG and Myc tags was also obtained for Exo70 (Fig. 5c and Figure S5B). MT labeling, however, appeared to be compromised using FLAG/Myc GCE-tags, with very few cells exhibiting MT staining using the Myc GCE-tag (Fig. 5c and Figure S5C). These results demonstrate the versatility of the system, in terms of the epitope used in the tag, and suggest that the GCE-tag can be employed both for fluorescent labeling and for other classical applications involving these epitopes.

**Fig 5.**
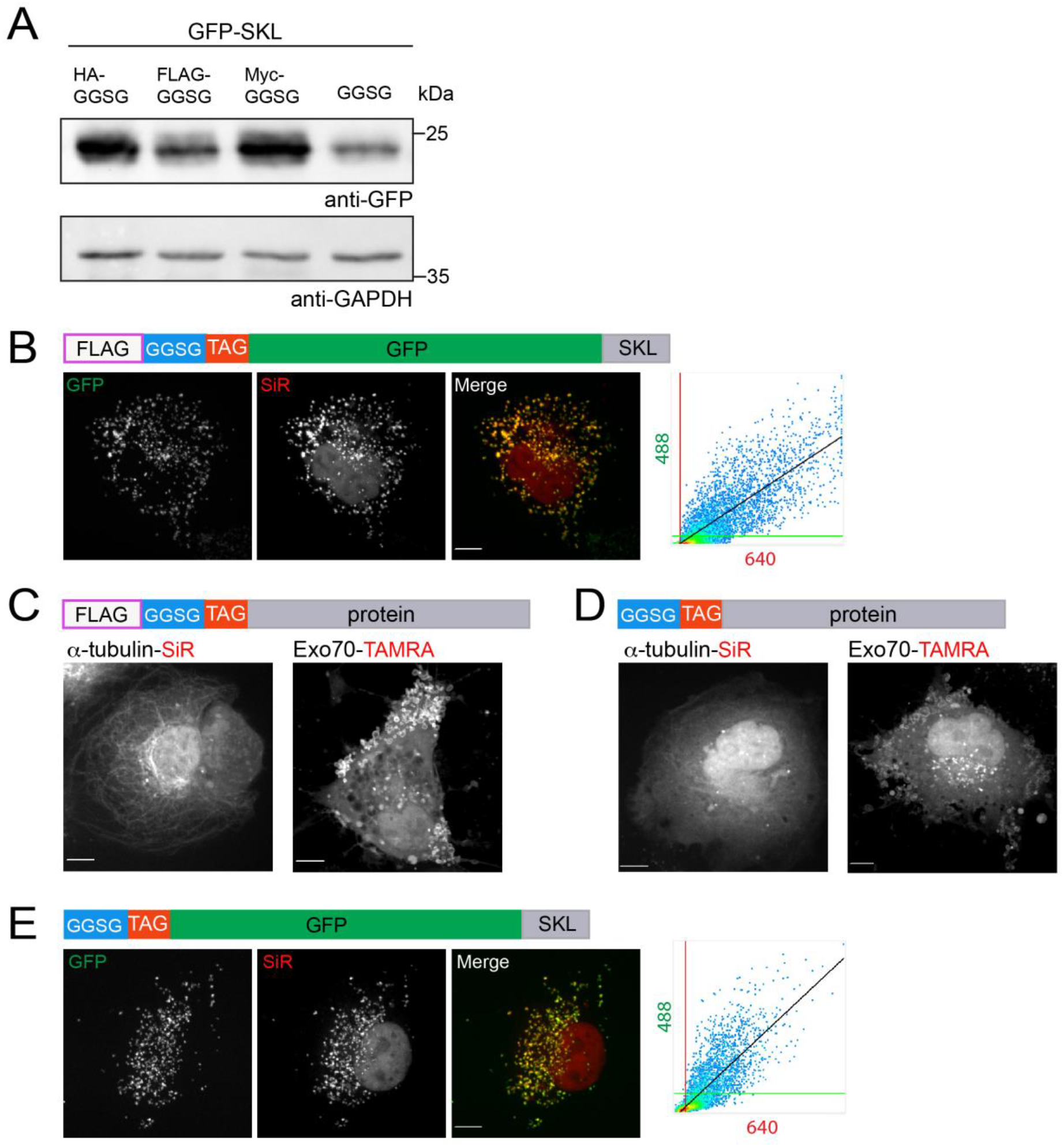
The GCE-tag is versatile and in some cases can be further minimized. (A) Western blot analysis of HEK293T cells transfected with pBUD-Pyl-RS plasmid inserted with GFP-SKL N-terminally tagged with (from left to right): the optimized GCE-tag (HA-GGSG-ncAA), tags with alternative short epitopes FLAG-GGSG-ncAA and Myc-GGSG-ncAA and a tag with no epitope (GGSG-ncAA). Cells were incubated for 48 h in the presence BCN-Lysine, harvested and subjected to western blot analysis using anti-GFP antibodies. (B-E) Live cell images of COS7 cells transfected as in (A) with FLAG-GGSG-ncAA (B and C), GGSG-ncAA (D and E) conjugated to: GFP-SKL (B and E), α-tubulin or Exo70 (C and D) and labeled with the indicated tetrazine dyes. Plots in (B and E) represent colocalization analysis between intensity values of the 488 (GFP) and 640 (SiR) channels performed on large subsets of the cells that excludes the nucleus, pearson correlation values: b, 0.858; c, 0.766. Results presented in each panel were obtained in at least 3 independent experiments. Scale-bar: 10 μm.

Last, we tested how removing the epitope sequence from the GCE-tag will affect labeling (Figure S1I). In other words, does a GGSG linker followed by BCN-Lys allow bioorthogonal labeling of proteins? Almost no MT labeling was obtained upon tagging α-tubulin with a GCE-tag that lacks the HA sequence, indicating that the residues contributed by the epitope are required for efficient labeling of MTs. For peroxisomes and exosomes, specific labeling was obtained, albeit at reduced levels (Fig. 5d, e, Pearson correlations in e, 0.766, respectively). These results indicate that in specific cases in which the length of tag is critical for preserving the function of the protein, the GCE-tag can be reduced to as few as 5 residues. Yet, this comes at the price of labeling efficiency and thus should be tested on a case-by-case basis.

## DISCUSSION

In this work we present a universal, small tag for labeling proteins in live mammalian cells with Fl-dyes through GCE and bioorthogonal chemistry, using a single expression vector. Efficient and specific labelling with Fl-dyes was observed for various intracellular structures and compartments, including MTs, PM, exosomes, lysosomes, MVBs, peroxisomes and ER using appropriate tag-bearing markers. By adding 14 residues to the N terminal of proteins we precluded the need for prior knowledge of the protein and bypassed the screening step currently associated with the technique. Moreover, by inserting the TAG codon at the beginning of the coding sequence (rather than in the middle of the sequence) we avoided the expression of truncated proteins. As labeling efficiency was not compensated by the use of the tag, the GCE-tag presented here provide an attractive, easy-to-implement, alternative for bioorthogonal labeling of proteins modified to carry a ncAA.

The GCE-tag reported here is considerably smaller than Fl-protein tags and self-labeling protein tags (GCE-tag, ~1.5 kDa; GFP, 27 kDa; SNAP-tag 19kDa) (1, 2). The only tag with a comparable size to that of the GCE-tag is the 12 amino acid long FlAsH tag (25). However, labeling proteins with FlAsH tags rely on biarsenical-functionalized fluorescent dyes, which exhibits unspecific biding to cellular membranes and are toxic to cells (2). Therefore, the GCE-tag stands as an attractive tool for labeling proteins and organelles in cells.

The library of GCE-tagged organelle-specific markers presented here is expected to be superior to currently available organelle markers. First, labeling is based on Fl-dyes, which are brighter and less sensitive to photobleaching than Fl-proteins. Second, the small size of the tag is likely to better preserve the physiological properties of the organelles, providing more realistic information on organellar dynamics in cells. Combining these markers with conventional organelle-specific markers or protein tags will allow for improved studies of organelle-organelle and organelle-protein interactions.

While demonstrating the suitability of the GCE-tag for labeling proteins in a variety of cellular compartments, the results described here stress the need to test several dyes when calibrating conditions for bioorthogonal labeling. This is mostly because organic dyes differ in their chemical properties and cellular compartments present different chemical environments. Table 1, summarizing the dyes found suitable for labeling each organelle, can be used as a guideline when optimizing labeling conditions for different cellular proteins. The increasing number of commercially available tetrazine-conjugated dyes will no doubt expand the ability to tailor Fl-dyes to specific cellular environments.

**Table 1.** List of organelle markers labeled via bioorthogonal labeling using the GCE-tag

| Organelle | Marker name | Fl-dye-Tet | Notes |
| --- | --- | --- | --- |
| Microtubules | α-tubulin | SiR | |
| Cell Membrane | CAAX | SiR/TAMRA | GFP Conjugated |
| Peroxisomes | SKL | SiR/TAMRA | GFP Conjugated |
| Lysosomes | Lamp1 | SiR |  |
| ER | ER<sup>cb5</sup>TM | TAMRA |  |
| Exosomes | Exo70 | TAMRA |  |
| MVBs | CD63 | TAMRA |  |

The use of the GCE-tag reported here can potentially be expanded to labeling endogenous proteins using genome-editing techniques and can be further applied to recently reported GCE-modified tissues and organisms (26–30). Additionally, a variety of ncAAs that carries different chemical functionalities have been genetically encoded in mammalian cells (8, 31-34). The GCE-tag can be further used for incorporating these ncAAs expanding its use in labeling applications and beyond.

## MATERIALS AND METHODS

### Cell culture

COS7 and HEK293T cells were grown in Dulbecco’s Modified Eagle Medium (DMEM; Life Technologies, Carlsbad, CA) supplemented with 10% fetal bovine serum (FBS), 2 mM glutamine, 10,000 U/ml penicillin and 10 mg/ml streptomycin.

### Plasmids and constructs

Tags were sub-cloned into the single expression vector pBUD-BCNK-RS that carries *pylT* encoding for tRNA_CUA_^Pyl^, and Pyrrolysyl-tRNA synthetase (18) (Figure S1), using *NotI/KpnI* restriction sites. Sequences encoding organelle markers were then inserted in frame using the *KpnI/XhoI* restriction sites at pBUD-BCNK-RS. All constructs were sequenced before use.

### Incorporation of the ncAA to proteins in cells

Assay was performed according to the our previously optimized protocol (6). Twenty-four hours prior to transfection, cells were plated at 20% confluency using the following dishes: live cell imaging, 4-well chamber slide (Ibidi, Martinsried, Germany); Western blot, 12-well plate (NUNC, Rochester, NY); immunostaining, #1.0 coverslips (Menzel, Braunschweig, Germany). Cells were transfected with pBUD-BCNK-RS plasmids carrying different tags and organelle markers (Figure S1 and Table S1) using Lipofectamine 2000 (Life Technologies, Carlsbad, CA) according to the manufacturer’s protocol, and incubated for 48 h in the presence of the ncAA BCN-Lysine (0.5 mM; Synaffix, Oss, Netherlands) in growth media supplemented with 100 μM ascorbic acid (Sigma Aldrich, Israel).

### Bioorthogonal labeling

48 h post transfection, cells were washed with fresh medium (3 × quick wash followed by 3 × 30 min wash) at 37°C, incubated with SiR-Tet (1–2 µM, 1 h; Spirochrome, Stein am Rhein, Switzerland) or TAMRA-Tet (2 µM, 1 h; Jena BioScience, Germany) and washed again with fresh medium (3 × quick wash and 3 × 30 min wash) at 37°C.

### Western Blot

Cells were harvested 48 h post transfection using RIPA lysis buffer (150 mM NaCl, 1% NP-40, 0.5% deoxycholate, 0.1% SDS, 50 mM Tris [pH 8.0]) supplemented with complete protease inhibitor for 30 min at 4°C. Total protein concentrations were measured with BCA Protein Assay Kit (Pierce Biotechnology) and equal total protein amounts were loaded. Membranes were stained with the following primary antibodies: rabbit anti-HA (1:4000; Applied Biological Materials, Richmond, Canada), mouse anti-GAPDH (1:1000; Applied Biological Materials), mouse anti-GFP (1:1000, Applied Biological Materials) and with rabbit or mouse-peroxidase secondary antibodies (1:10,000; Jackson ImmunoResearch, West Grove, PA).

### Immunofluorescence

Cells were fixed 48 h post transfection with 4% paraformaldehyde (PFA) and co-stained with rabbit anti-HA (1:500) and Mouse anti-CD63 (1:200; Abcam, Cambridge, MA) primary antibodies and with Alexa Fluor 488 anti-rabbit and Alexa Fluor 594 anti-mouse (1:500, Life Technologies) secondary antibodies. Cells were mounted with Fluoromount-G (SouthernBiotech, Birmingham, AL).

### Live cell imaging and image processing

Cells were imaged on a fully incubated confocal spinning-disk microscope at 37°C (Marianas; Intelligent Imaging, Denver, CO) using a 63× oil objective (numerical aperture 1.4), and recorded on an electron-multiplying charge-coupled device camera (pixel size, 0.079 μm; Evolve; Photometrics, Tucson, AZ). FRAP experiments were performed on cells expressing GCE-tag-ER^cb5^TM and labeled with TAMRA-Tet. After 4 baseline time-points, bleaching was carried out using a 405 laser. Recovery after bleaching was recorded for 1 min with 1.8 s intervals. Image processing, FRAP analysis fitting and unintentional bleaching corrections were performed using SlideBook, version 6 (Intelligent Imaging, Denver, CO).

### In vitro Analysis of pH sensitivity of Tetrazine conjugated Fl-dyes

SiR-Tet (1µM) or TAMRA-Tet (2 µM) dyes were diluted in HEPES buffers of different acidity (pH ranges from 5 to 9) in the presence or absence of the ncAA BCN-Lysine (0.5 mM). As a control, Fl-dyes were also diluted in 0.1 M HCl and 0.1 M NaOH. Samples were then loaded in triplicates onto a 384-well plate (Greiner), and incubated for 30 min at 37°C. Fluorescence intensity was recorded using infinite M1000 plate reader (Tecan, Männedorf, Switzerland) using 652/674 nm or 545/575 nm excitation / emission wavelengths, respectively.

## ACKNOWLEDGMENTS

We thank Peter Kim (university of Toronto) and Jennifer Lippincott Schwartz (Janelia Research campus) for contributing plasmids for fluorescent protein based organelle markers. We also thank all members of the Elia lab for critical reading of the manuscript. The project leading to this application has received funding from European Research Council (ERC) under the European Union’s Horizon 2020 research.

## AUTHOR CONTRIBUTIONS

I.S and N.E designed and analyzed all the experiments. I.S performed all the experiments. D.N provided technical support. E.A advised on GCE. N.E, EA and DN initiated the project. N.E wrote the manuscript.

---

**Figure S1.**
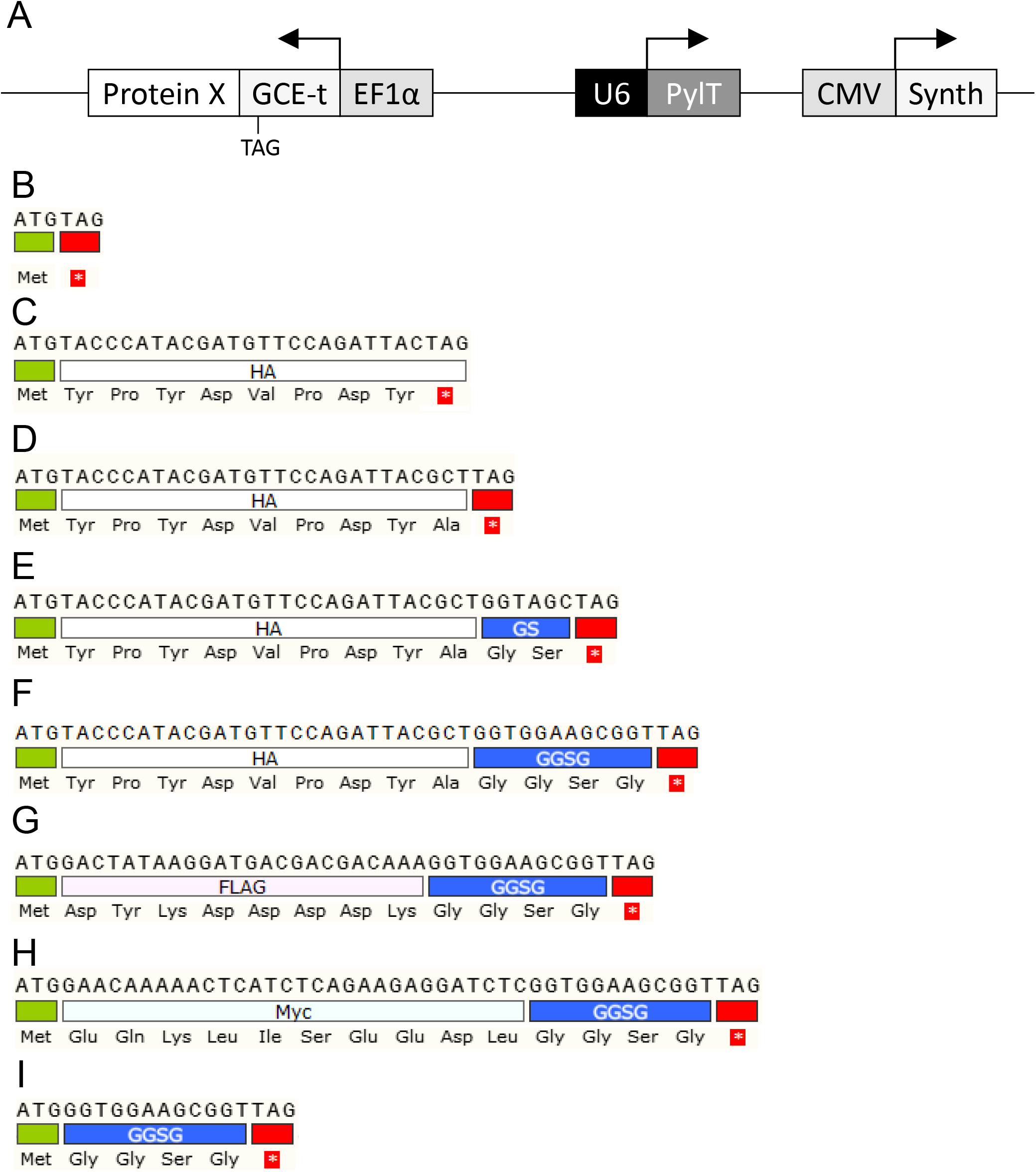
Single expression vector maps and sequences of the designed linkers. (A) Schematic representation of the genetic code expansion vector with *PylT* gene and BCN-RS. (B-I) Nucleotide sequences of (B) Methionine-TAG, (C) HA*-TAG (corresponds to tag 1 in Fig. 1 C), (D) HA-TAG (corresponds to tag 2 in Fig. 1 C), (E) HA-GS-TAG (corresponds to tag 3 in Fig. 1 C), (F) HA-GGSG-TAG (corresponds to tag 4 in Fig. 1 C), (G) Flag-GGSG-TAG, (H) Myc-GGSG-TAG, (I) GGSG-TAG. ATG represents beginning of ORF, ncAA incorporation site is at the TAG codon marked in red.

**Figure S2.**
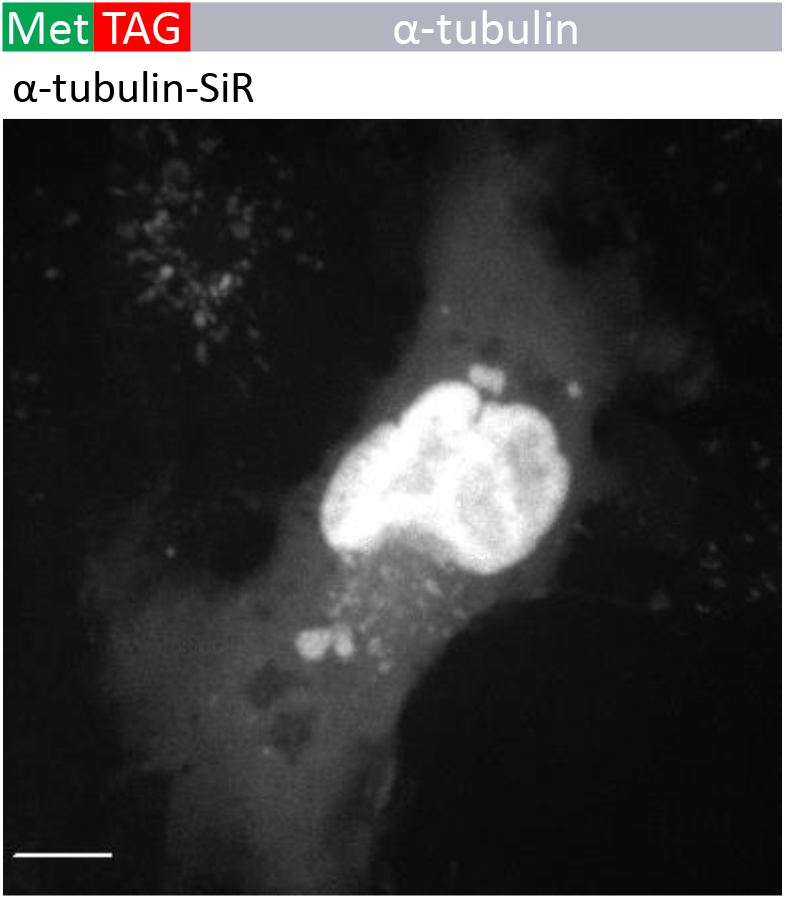
Negative MT labeling with Met-TAG-α-tubulin. COS7 cells were transfected with pBUD-Pyl-RS that carries Methionine-TAG-α-tubulin, and labeled with SiR-Tet as described in the materials and methods. Note that there was no MT labeling using this Tag. Scale-bar: 10 μm

**Figure S3.**
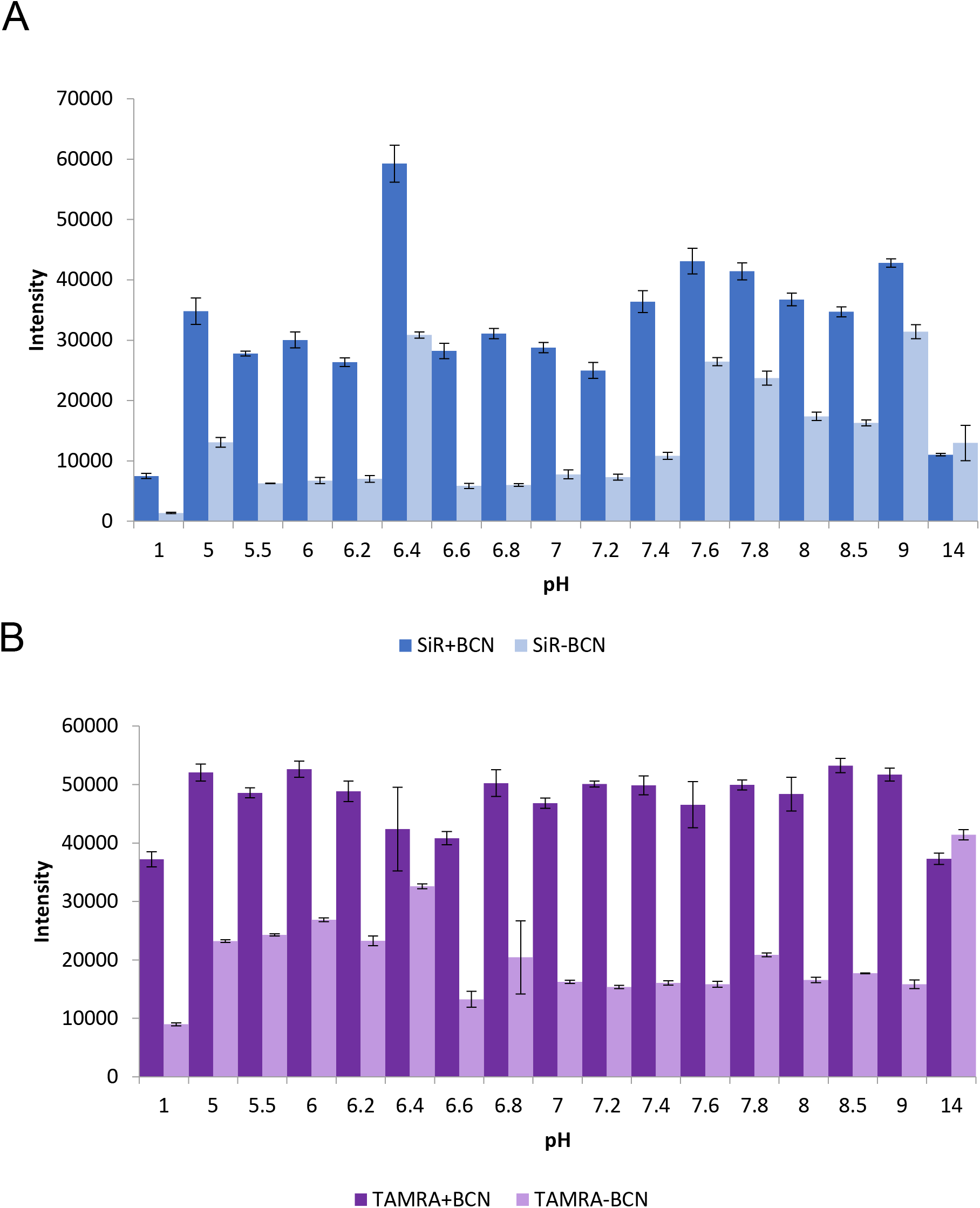
pH sensitivity of Tetrazine conjugated Fl-dyes. In vitro analysis of SiR-Tet (A) and TAMRA-Tet (B) diluted in HEPES buffer in different pH, in the presence or absence of the ncAA BCN-Lysine as described in the supplementary methods (means ± s.d.)

**Figure S4.**
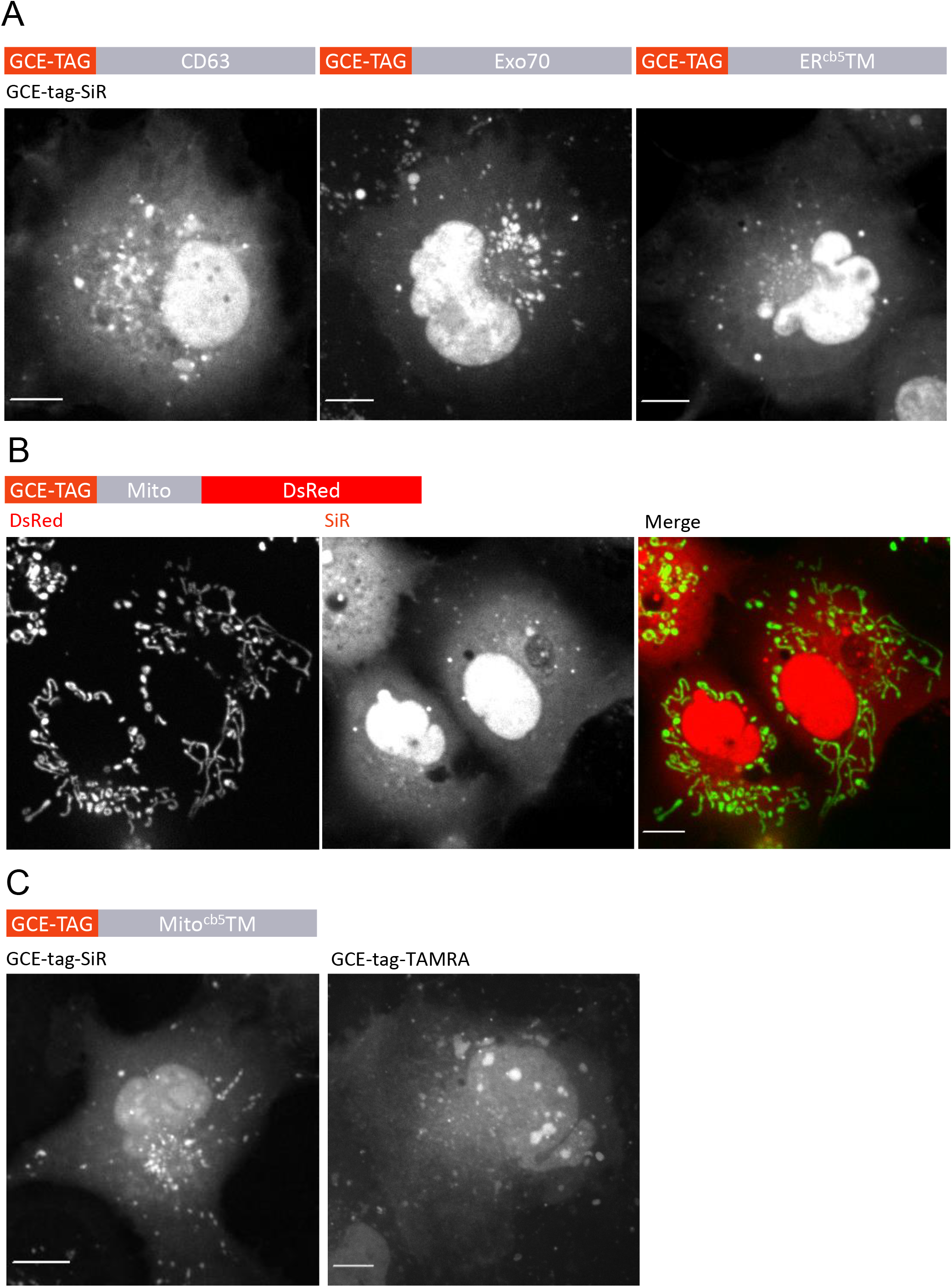
Negative labeling of mitochondria, MVBs, Exosomes and ER using the GCE-tag. COS7 cells transfected with pBUD-Pyl-RS carrying (A) from left to right: GCE-tag-CD63, GCE-tag-Exo70 or GCE-tag-ER^cb5^TM, (B) GCE-tag-Mito-DsRed, or (C) GCE-tag-Mito^cb5^TM. Cells were labeled with SiR-Tet (A and B) or TAMRA-Tet (C), as described in the materials and methods. No specific labeling was obtained under any of these conditions. Scale-bar: 10 μm.

**Figure S5.**
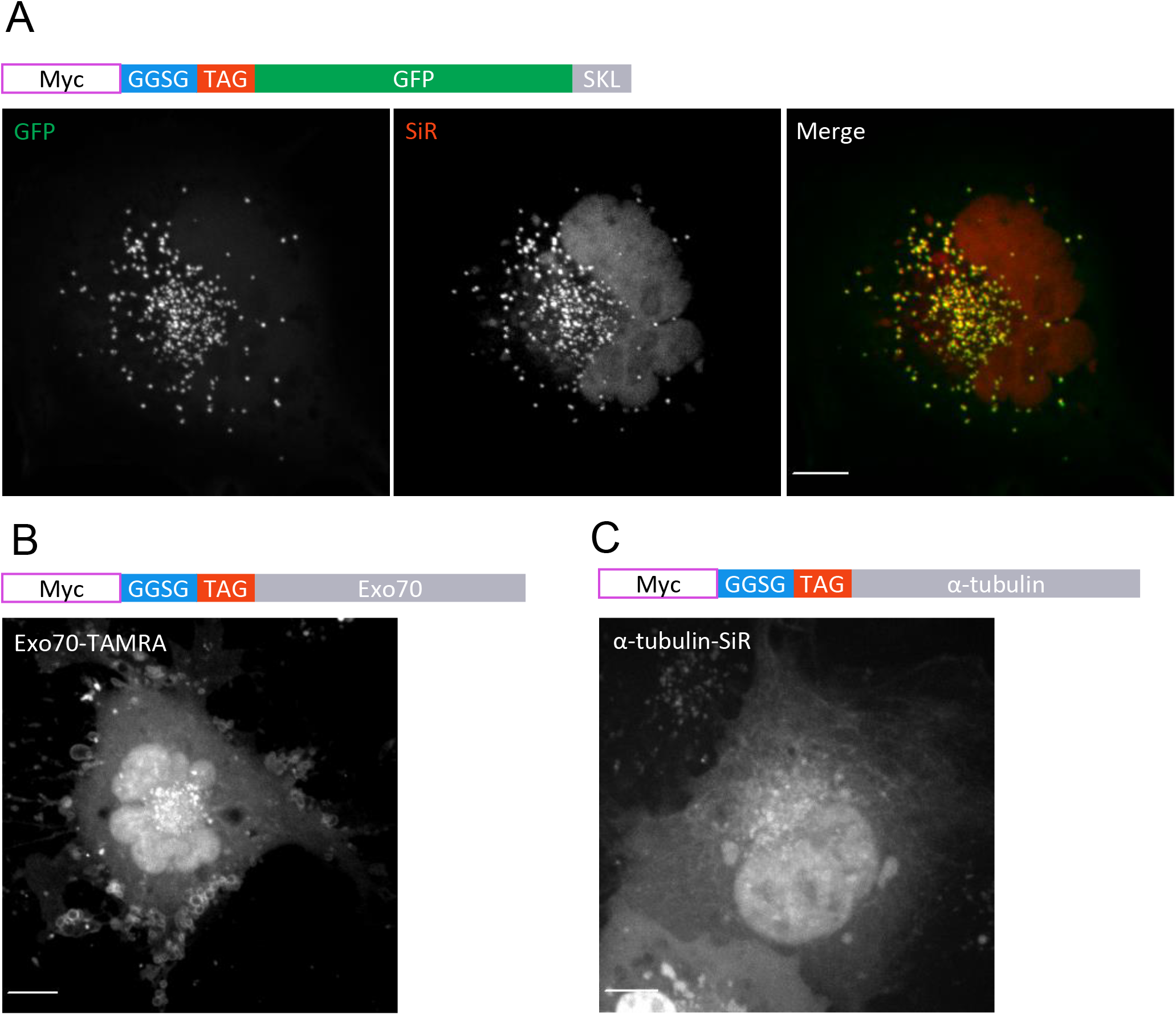
Peroxisome, Exosome and MT labeling using Myc-GGSG-TAG as a bioorthogonal labeling tag. COS7 cells transfected with pBUD-Pyl-RS carrying either SKL-GFP (A), Exo70 (B), or α-tubulin (C), all conjugated to the tag Myc-GGSG-TAG at the N-terminus, and labeled with (A and C) SiR-Tet or (B) TAMRA-Tet as described in the materials and methods, indicating that in some target proteins the HA epitope can be replaced by Myc. Scale-bar: 10 μm.

**Table S1.**
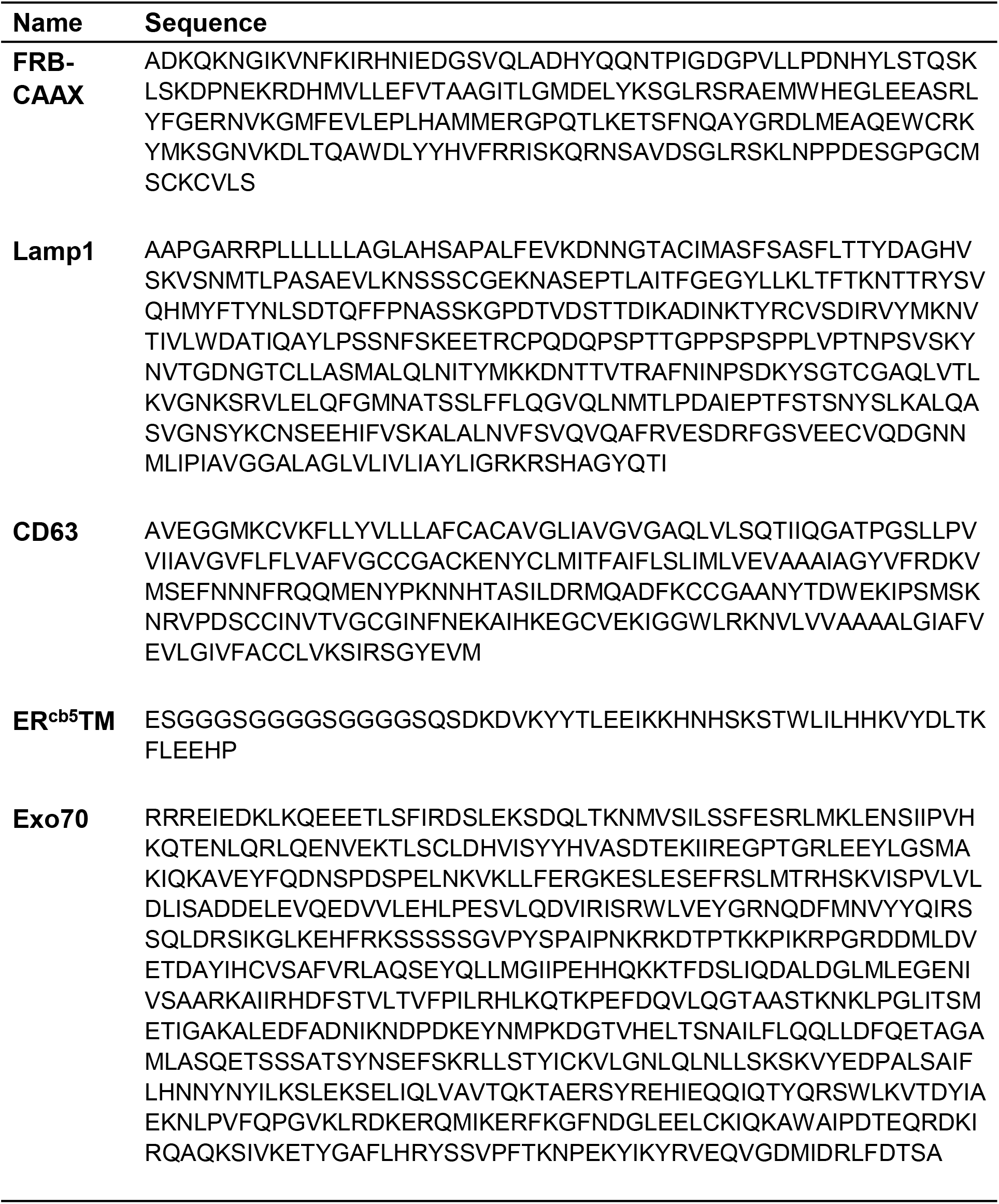
Sequences of organelle markers used in this work

