## Supplemental Figures, Tables and Videos for "A straightforward approach for bioorthogonal labeling of proteins and organelles in live mammalian cells, using a short peptide tag"

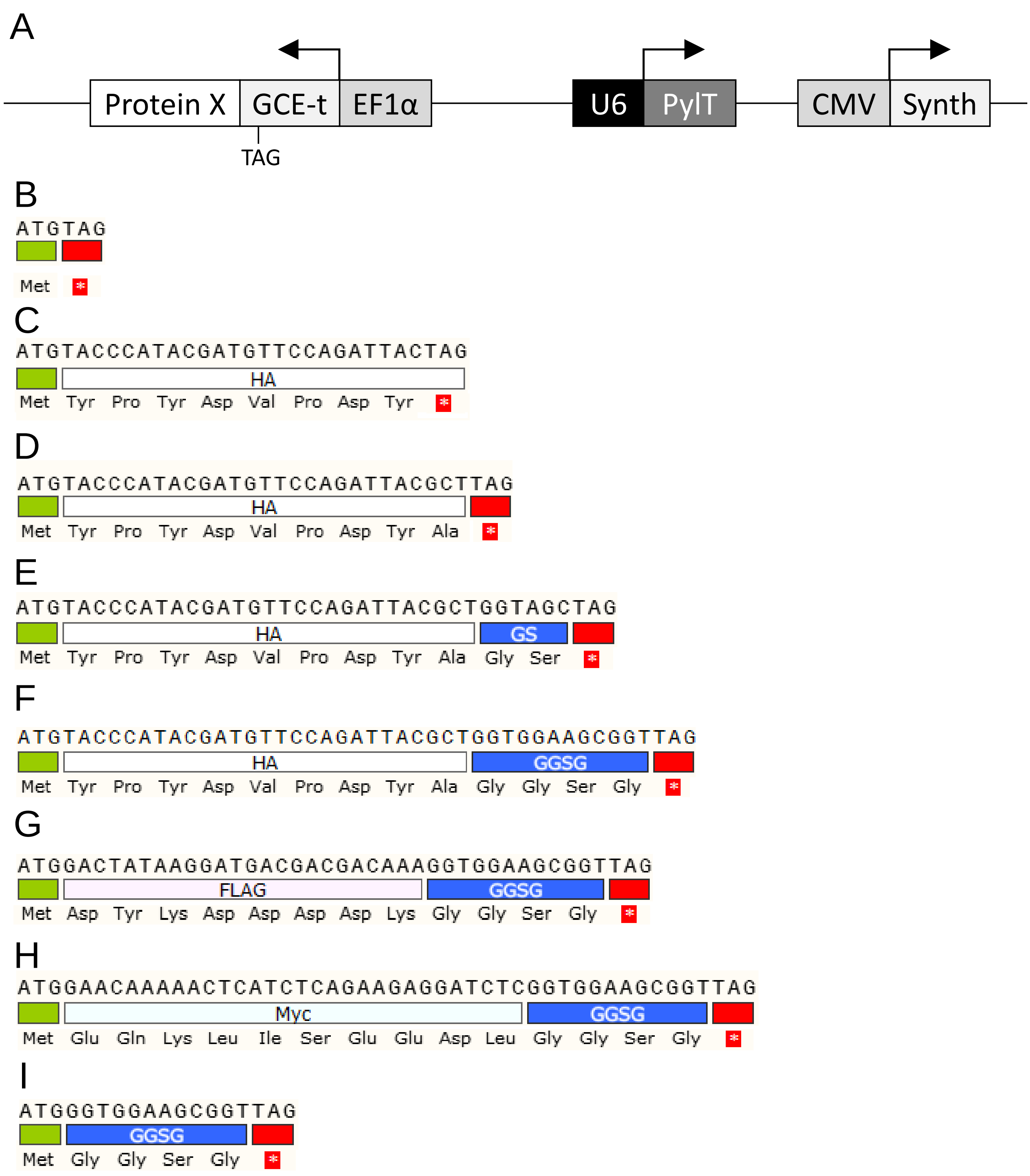


**Figure S1. Single expression vector maps and sequences of the designed linkers**
*Related to Figure 2*

(A) Schematic representation of the genetic code expansion vector with *PylT* gene and BCN-RS. (B-I) Nucleotide sequences of (B) Methionine-TAG, (C) HA*-TAG (corresponds to tag 1 in Fig. 1 C), (D) HA-TAG (corresponds to tag 2 in Fig. 1 C), (E) HA-GS-TAG (corresponds to tag 3 in Fig. 1 C), (F) HA-GGSG-TAG (corresponds to tag 4 in Fig. 1 C), (G) Flag-GGSG-TAG, (H) Myc-GGSG-TAG, (I) GGSG-TAG. ATG represents beginning of ORF, ncAA incorporation site is at the TAG codon marked in red.

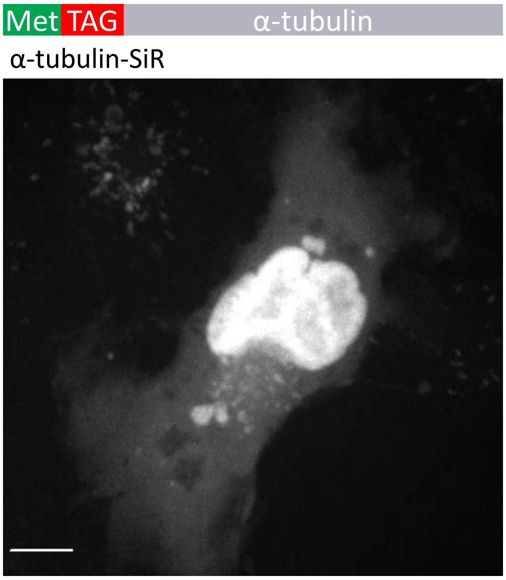


**Figure S2. Negative MT labeling with Met-TAG-α-tubulin**
*Related to Figure 2*

COS7 cells were transfected with pBUD-Pyl-RS that carries Methionine-TAG-α-tubulin, and labeled with SiR-Tet as described in the materials and methods. Note that there was no MT labeling using this Tag. Scale-bar: 10 μm

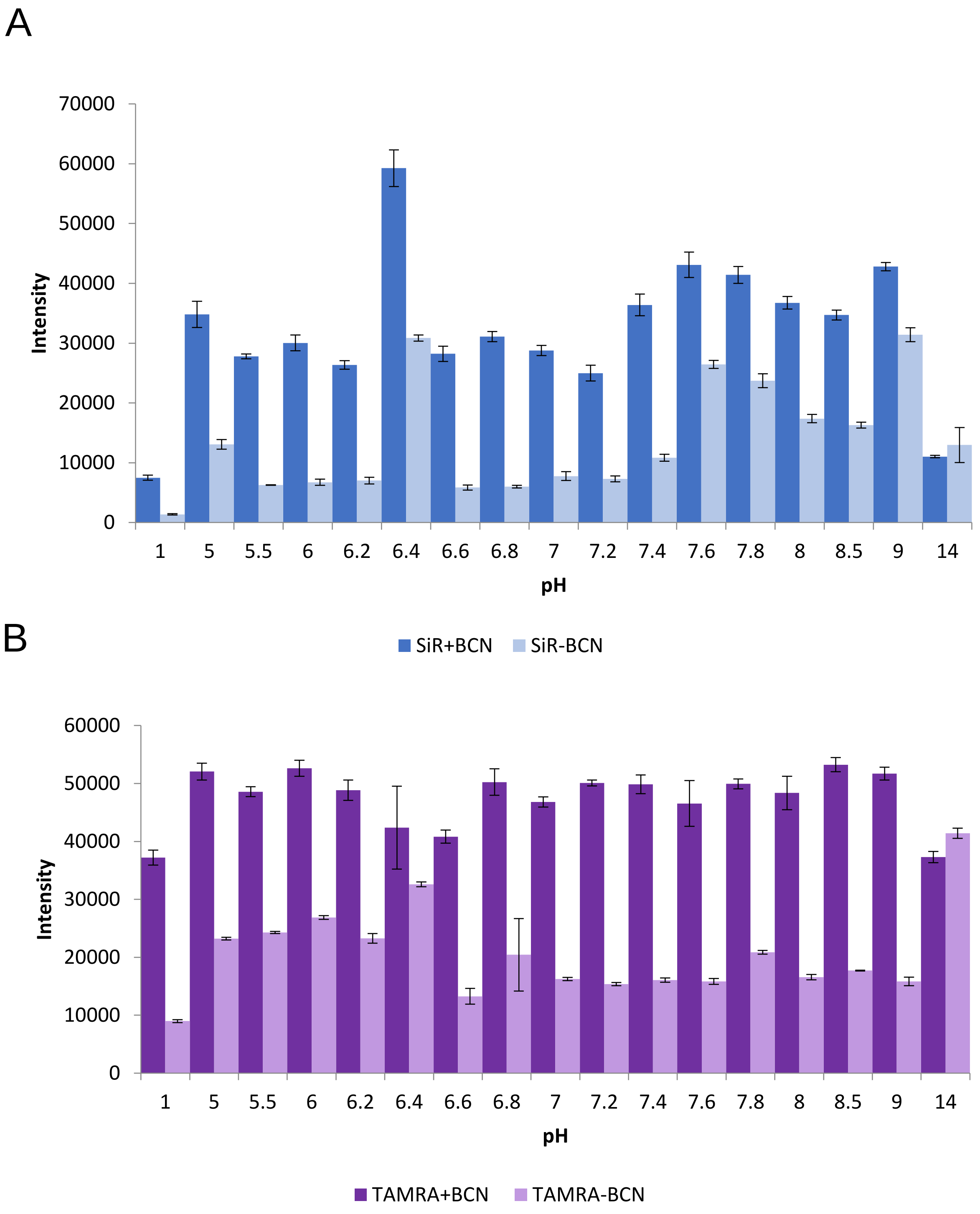


**Figure S3. pH sensitivity of Tetrazine conjugated Fl-dyes**
*Related to Figure 3*

In vitro analysis of SiR-Tet (A) and TAMRA-Tet (B) diluted in HEPES buffer in different pH, in the presence or absence of the ncAA BCN-Lysine as described in the supplementary methods (means ± s.d.)

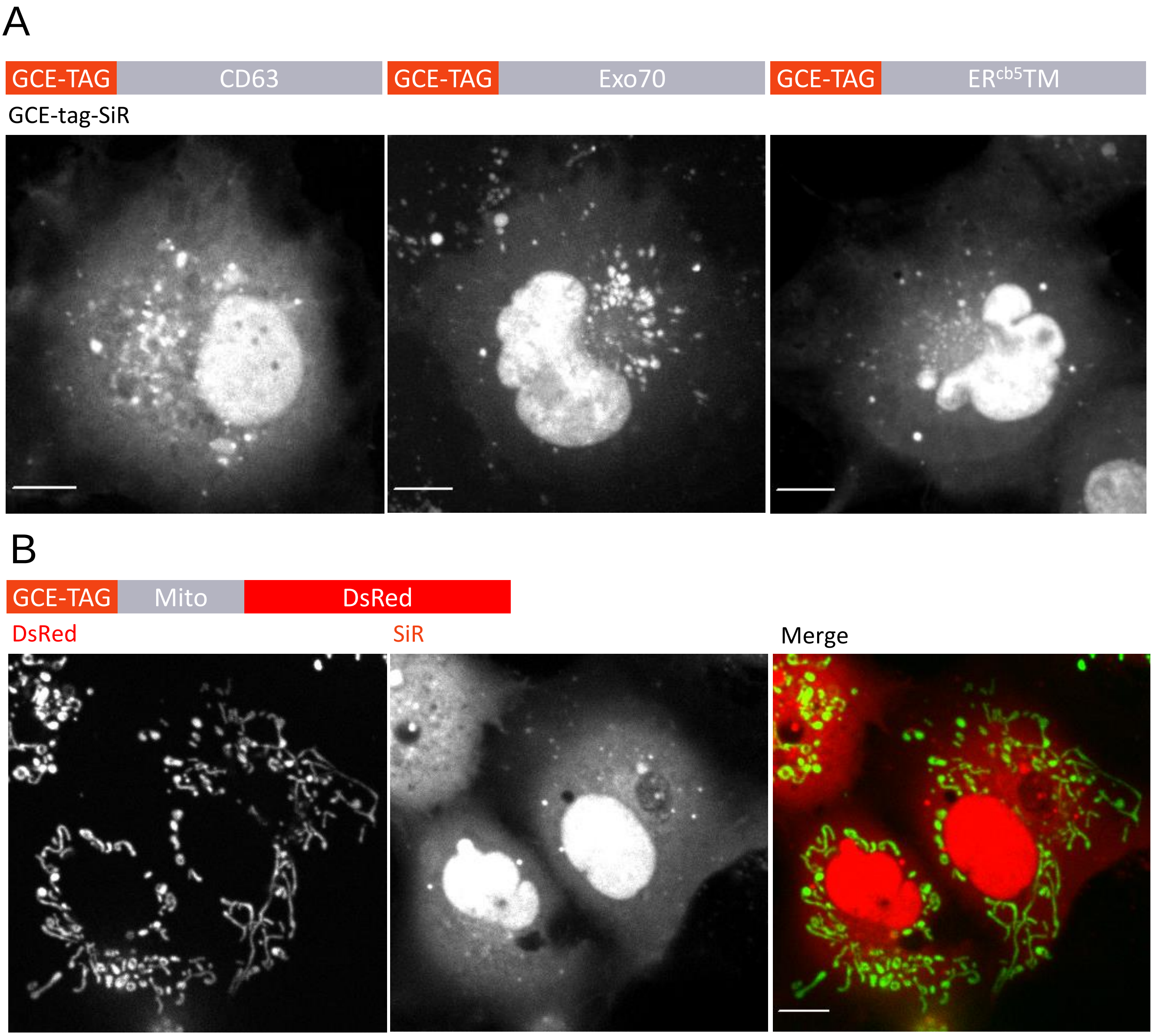


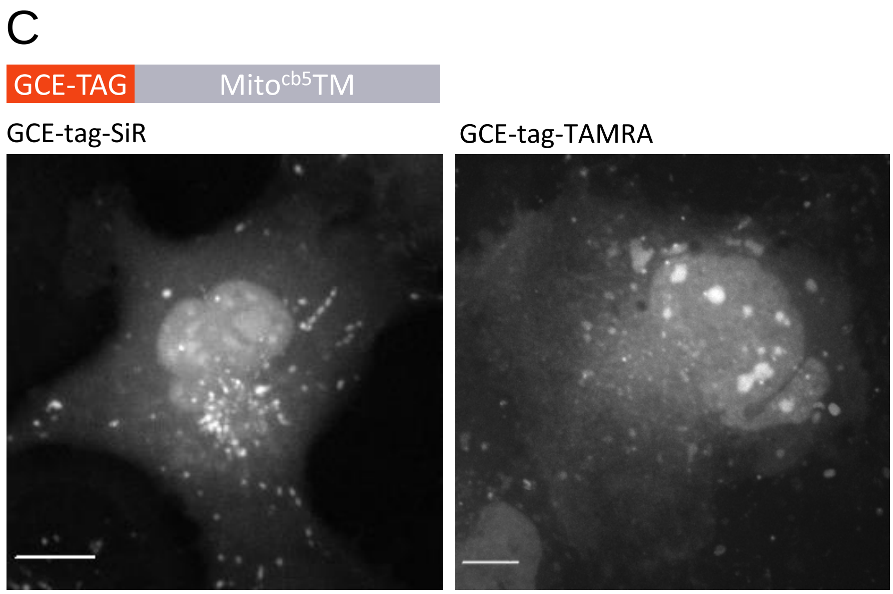


**Figure S4. Negative labeling of mitochondria, MVBs, Exosomes and ER using the GCE-tag.**
*Related to Figure 4*

COS7 cells transfected with pBUD-Pyl-RS carrying (A) from left to right: GCE-tag-CD63, GCE-tag-Exo70 or GCE-tag-ER$^{cb5}$TM, (B) GCE-tag-Mito-DsRed, or (C) GCE-tag-Mito$^{cb5}$TM. Cells were labeled with SiR-Tet (A and B) or TAMRA-Tet (C), as described in the materials and methods. No specific labeling was obtained under any of these conditions. Scale-bar: 10 μm.

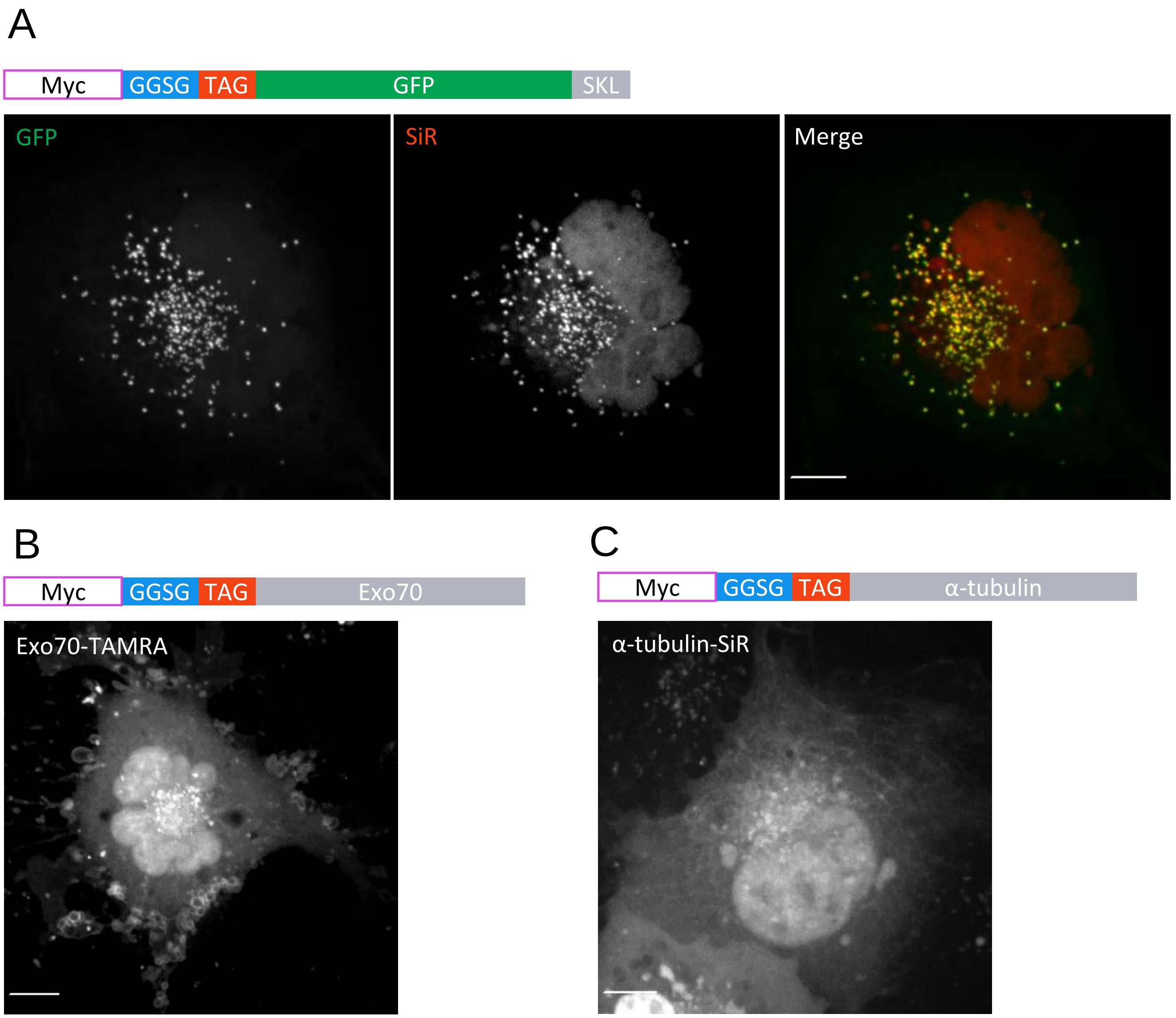


**Figure S5. Peroxisome, Exosome and MT labeling using Myc-GGSG-TAG as a bioorthogonal labeling tag**
*Related to Figure 5*

COS7 cells transfected with pBUD-Pyl-RS carrying either SKL-GFP (A), Exo70 (B), or α-tubulin (C), all conjugated to the tag Myc-GGSG-TAG at the N-terminus, and labeled with (A and C) SiR-Tet or (B) TAMRA-Tet as described in the materials and methods, indicating that in some target proteins the HA epitope can be replaced by Myc. Scale-bar: 10 µm.

| Name | Sequence |
|---|---|
| **FRB-CAAX** | ADKQKNGIKVNFKIRHNIEDGSVQLADHYQQNTPIGDGPVLLPDNHYLSTQSK LSKDPNEKRDHMVLLEFVTAAGITLGMDELYKSGLRSRAEMWHEGLEEASRL YFGERNVKGMFEVLEPLHAMMERGPQTLKETSFNQAYGRDLMEAQEWCRK YMKSGNVKDLTQAWDLYYHVFRRISKQRNSAVDSGLRSKLNPPDESGPGCM SCKCVLS |
| **Lamp1** | AAPGARRPLLLLLLAGLAHSAPALFEVKDNNGTACIMASFSASFLTTYDAGHV SKVSNMTLPASAEVLKNSSSCGEKNASEPTLAITFGEGYLLKLTFTKNTTRYSV QHMYFTYNLSDTQFFPNASSKGPDTVDSTTDIKADINKTYRCVSDIRVYMKNV TIVLWDATIQAYLPSSNFSKEETRCPQDQPSPTTGPPSPSPPLVPTNPSVSKY NVTGDNGTCLLASMALQLNITYMKKDNTTVTRAFNINPSDKYSGTCGAQLVTL KVGNKSRVLELQFGMNATSSLFFLQGVQLNMTLPDAIEPTFSTSNYSLKALQA SVGNSYKCNSEEHIFVSKALALNVFSVQVQAFRVESDRFGSVEECVQDGNN MLIPIAVGGALAGLVLIVLIAYLIGRKRSHAGYQTI |
| **CD63** | AVEGGMKCVKFLLYVLLLAFCACAVGLIAVGVGAQLVLSQTIIQGATPGSLLPV VIIAVGVFLFLVAFVGCCGACKENYCLMITFAIFLSLIMLVEVAAAIAGYVFRDKV MSEFNNNFRQQMENYPKNNHTASILDRMQADFKCCGAANYTDWEKIPSMSK NRVPDSCCINVTVGCGINFNEKAIHKEGCVEKIGGWLRKNVLVVAAAALGIAFV EVLGIVFACCLVKSIRSGYEVM |
| **ER$^{cb5}$TM** | ESGGGSGGGGSGGGGSQSDKDVKYYTLEEIKKHNHSKSTWLILHHKVYDLTK FLEEHP |
| **Exo70** | RRREIEDKLKQEEETLSFIRDSLEKSDQLTKNMVSILSSFESRLMKLENSIIPVH KQTENLQRLQENVEKTLSCLDHVISYYHVASDTEKIIREGPTGRLEEYLGSMA KIQKAVEYFQDNSPDSPELNKVKLLFERGKESLESEFRSLMTRHSKVISPVLVL DLISADDELEVQEDVVLEHLPESVLQDVIRISRWLVEYGRNQDFMNVYYQIRS SQLDRSIKGLKEHFRKSSSSSGVPYSPAIPNKRKDTPTKKPIKRPGRDDMLDV ETDAYIHCVSAFVRLAQSEYQLLMGIIPEHHQKKTFDSLIQDALDGLMLEGENI VSAARKAIIRHDFSTVLTVFPILRHLKQTKPEFDQVLQGTAASTKNKLPGLITSM ETIGAKALEDFADNIKNDPDKEYNMPKDGTVHELTSNAILFLQQLLDFQETAGA MLASQETSSSATSYNSEFSKRLLSTYICKVLGNLQLNLLSKSKVYEDPALSAIF LHNNYNYILKSLEKSELIQLVAVTQKTAERSYREHIEQQIQTYQRSWLKVTDYIA EKNLPVFQPGVKLRDKERQMIKERFKGFNDGLEELCKIQKAWAIPDTEQRDKI RQAQKSIVKETYGAFLHRYSSVPFTKNPEKYIKYRVEQVGDMIDRLFDTSA |

**Table S1. Sequences of organelle markers used in this work**
*Related to Figure 3 and Table 1*

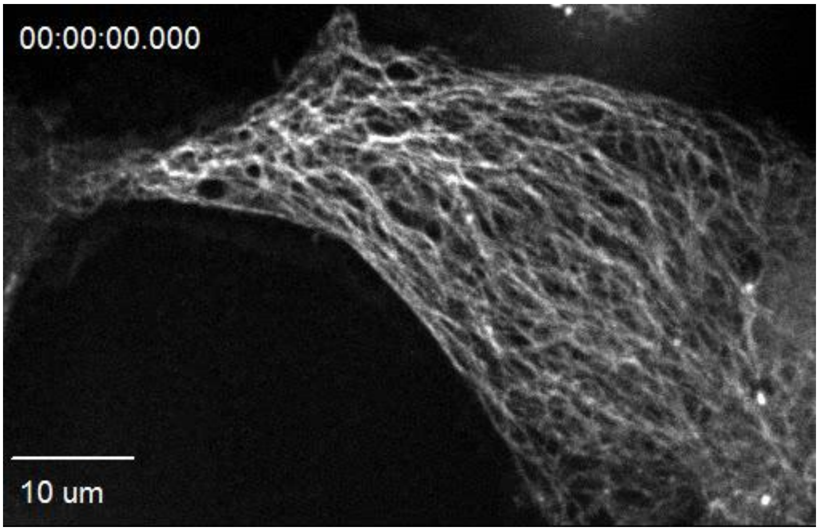


**Video S1.**
Microtubule dynamics recorded in COS7 cells expressing GCE-tag-α-tubulin and labeled with SiR-Tet. Cells were recorded for approximately 1 min with 4 s intervals. Scale-bar: 10 μm.

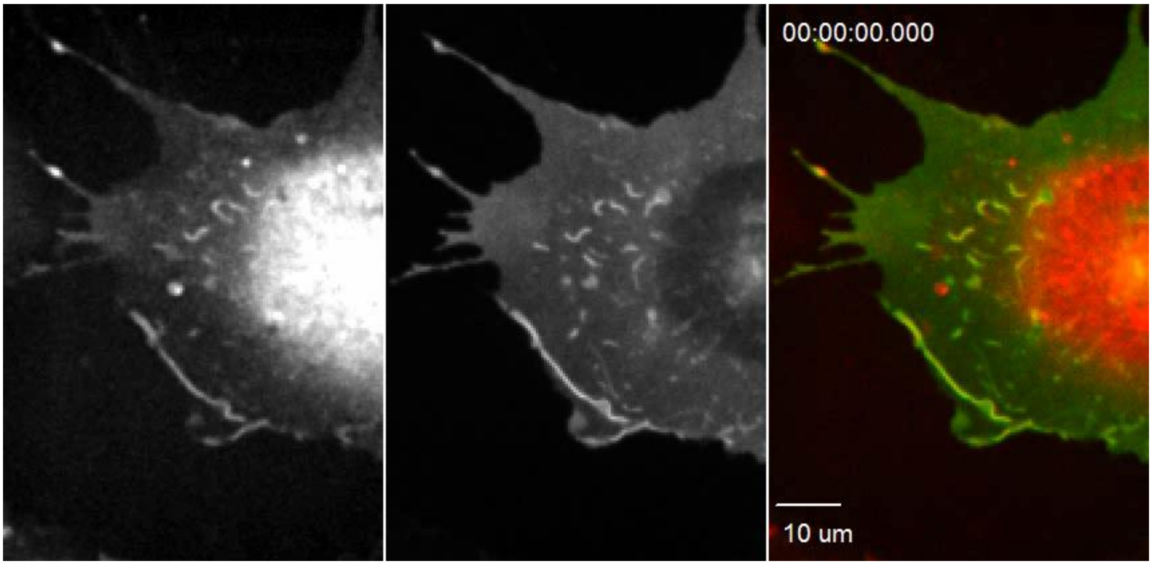


**Video S2.**
To induce plasma membrane dynamics, COS7 cells expressing GCE-tag-GFP-CAAX labeled with SiR-Tet, were incubated in serum-free DMEM for 6h and then washed with DMEM. Cells were recorded for 5 min with 5.5s intervals. Left panel: 640 (SiR, red) channel, middle panel: 488 (GFP, green) channel. Scale-bar: 10 μm.

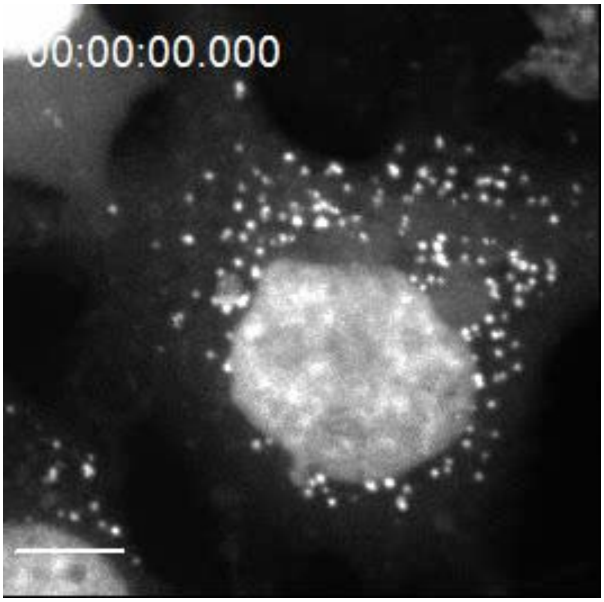


**Video S3.**
Peroxisome dynamics recorded in COS7 cells expressing GCE-tag-GFP-SKL and labeled with SiR-Tet. Cells were recorded for 5 min with 11 s intervals. Only SiR channel is shown. Scale-bar: 10 μm.

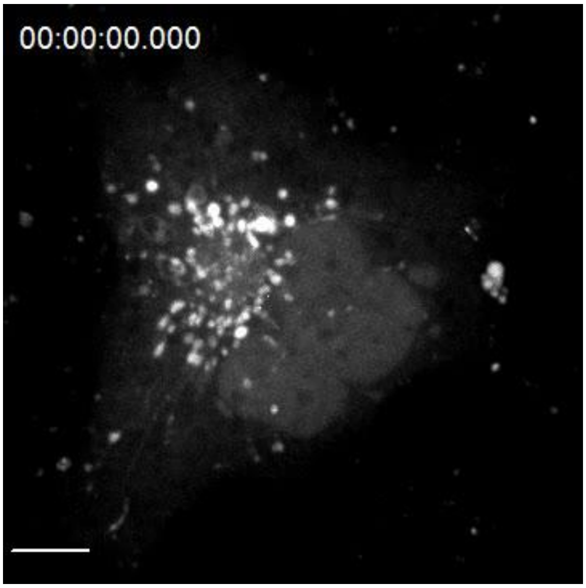


**Video S4.**
MVB dynamics recorded in COS7 cells expressing GCE-tag-CD63 and labeled with TAMRA-Tet. Cells were recorded for 5 min with 4 s intervals. Scale-bar: 10 μm.

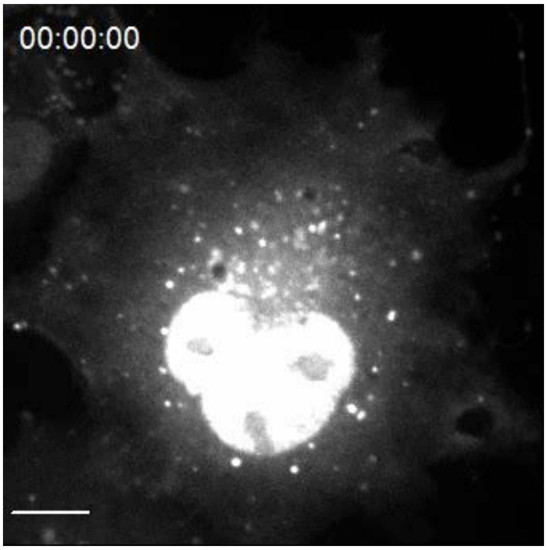


**Video S5.**
Lysosome inhibition recorded in COS7 cells expressing GCE-tag-Lamp1 and labeled with SiR-Tet. Cells were imaged for 3 h in the presence of the lysosome inhibitor chloroquine (120 μM), at 10 min intervals. Scale-bar: 10 μm.

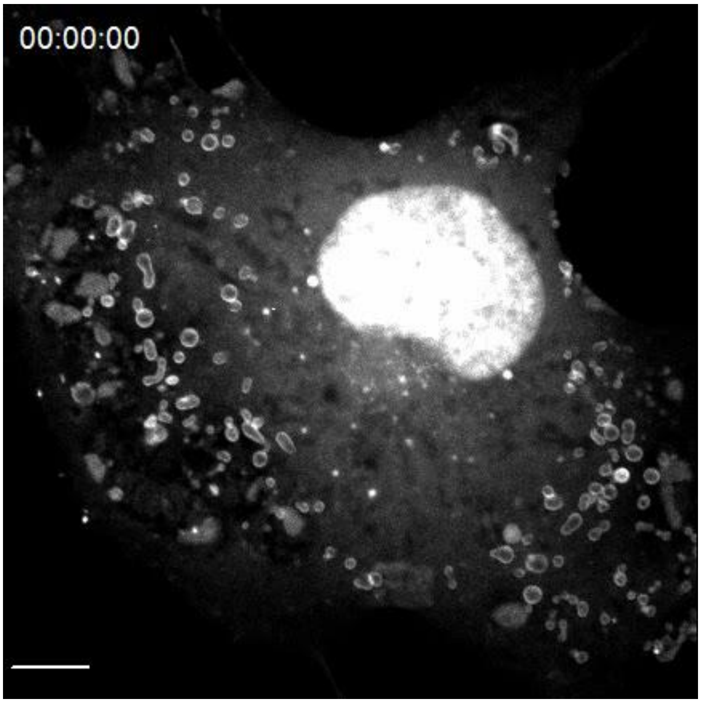


**Video S6.**
Exosomes dynamics recorded in COS7 cells expressing GCE-tag-Exo70 and labeled with TAMRA-Tet. Cells were recorded for 30 s with 1 s intervals. Scale-bar: 10 μm.

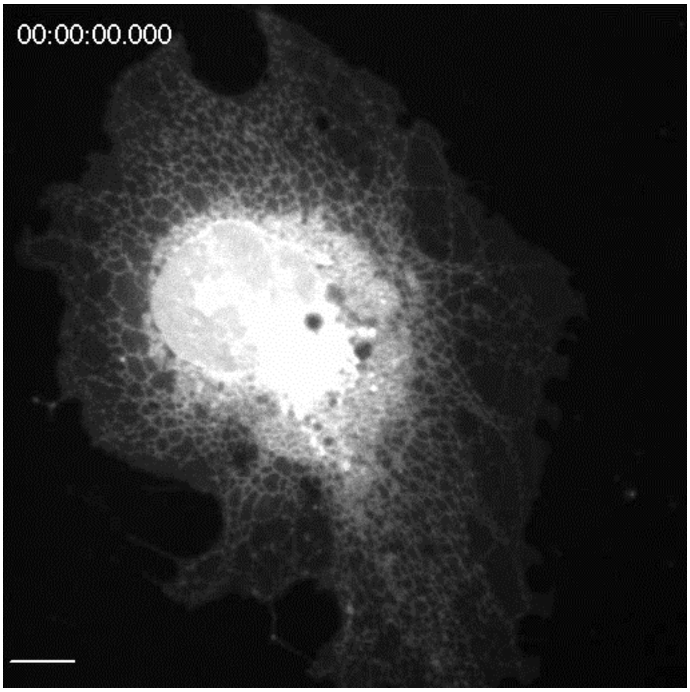


**Video S7.**
FRAP analysis of COS7 cells expressing GCE-tag-ER$^{cb5}$TM labeled with TAMTA-Tet. Cells were imaged for 2 min with 2 s intervals. Photo-bleaching was performed after 4 baseline timepoints (12 s). Scale-bar: 10 µm.

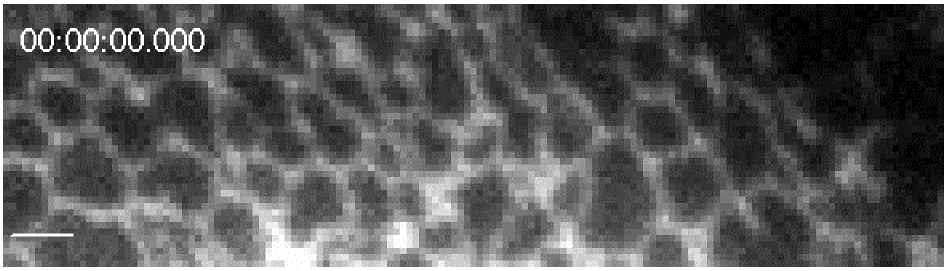


**Video S8.**
Zoomed-in video of the bleached ROI in cells expressing GCE-tag-ER$^{cb5}$TM labeled with TAMTA-Tet (taken from supplementary video 7). Cells were imaged for 2 min with 2 s intervals. Photo-bleaching was performed after 4 baseline timepoints (12 s). Scale-bar: 2 µm.
